# Photothermal Recycling Biosensing for Continuous, Sensitive Molecular Quantification

**DOI:** 10.64898/2026.03.30.714774

**Authors:** Yongchen Tai, Yunshen Li, Wenting Wang, Yang Lu, Ziyan Qian, Mitchell Conover, Josef Neu, Carl Denard, Qiye Zheng, Jing Pan

## Abstract

Continuous biochemical sensing provides valuable insights into an individual’s physiological state and the mechanisms underlying pathophysiological changes. However, most existing bioanalytical methods are not compatible with continuous biochemical sensing. A major technical challenge lies in achieving rapid measurement readouts while maintaining high specificity and sensitivity in complex biological fluids. Sensitive molecular detection typically requires slow analyte–binder dissociation and long incubation to reach equilibrium, whereas rapid and frequent measurements demand fast association–dissociation kinetics that are difficult to reconcile for low-abundance analytes. To address this challenge, we introduce a sensing mechanism termed photothermal recycling (PTR), which mimics the thermal cycling process in polymerase chain reaction. Using plasmonic photothermal effects, PTR rapidly recycles binders to enable frequent measurements. We demonstrate a digital PTR assay capable of multi-hour biochemical monitoring with subpicomolar(pM) sensitivity in buffer, diluted serum, and saliva. This approach leverages localized thermal energy to dynamically modulate biomolecular recognition, offering a new bioanalytical paradigm for continuous biochemical sensing across diverse application settings.

## 1. Introduction

Continuous monitoring that captures longitudinal biochemical dynamics provides valuable insights into an individual’s physiological state and overall health trajectory, enabling informed clinical decision-making and timely interventions.[1] Various continuous sensors have been previously reported, implementing different modalities such as electrochemical sensors, tethered particle assays, flow-based immunoassay, etc.[2–11] Most of the current continuous biosensors detect analytes ranging from picomolar(pM) to micromolar(µM) concentrations. The assay formats commonly used in these biosensors rely on analyte-binder interactions under equilibrium conditions. To achieve high sensitivity towards low-abundance analytes, biosensors often employ strong binders with high affinity and a low dissociation constant (Kd). However, once the binder-analyte complex is formed, the separation process is very slow, sacrificing the temporal response during continuous operations. Thus, biosensing mechanisms relying on equilibrium binding face limitations in achieving sensitivity and speed simultaneously in continuous biochemical monitoring applications.(Fig.1a)

**Fig. 1:**
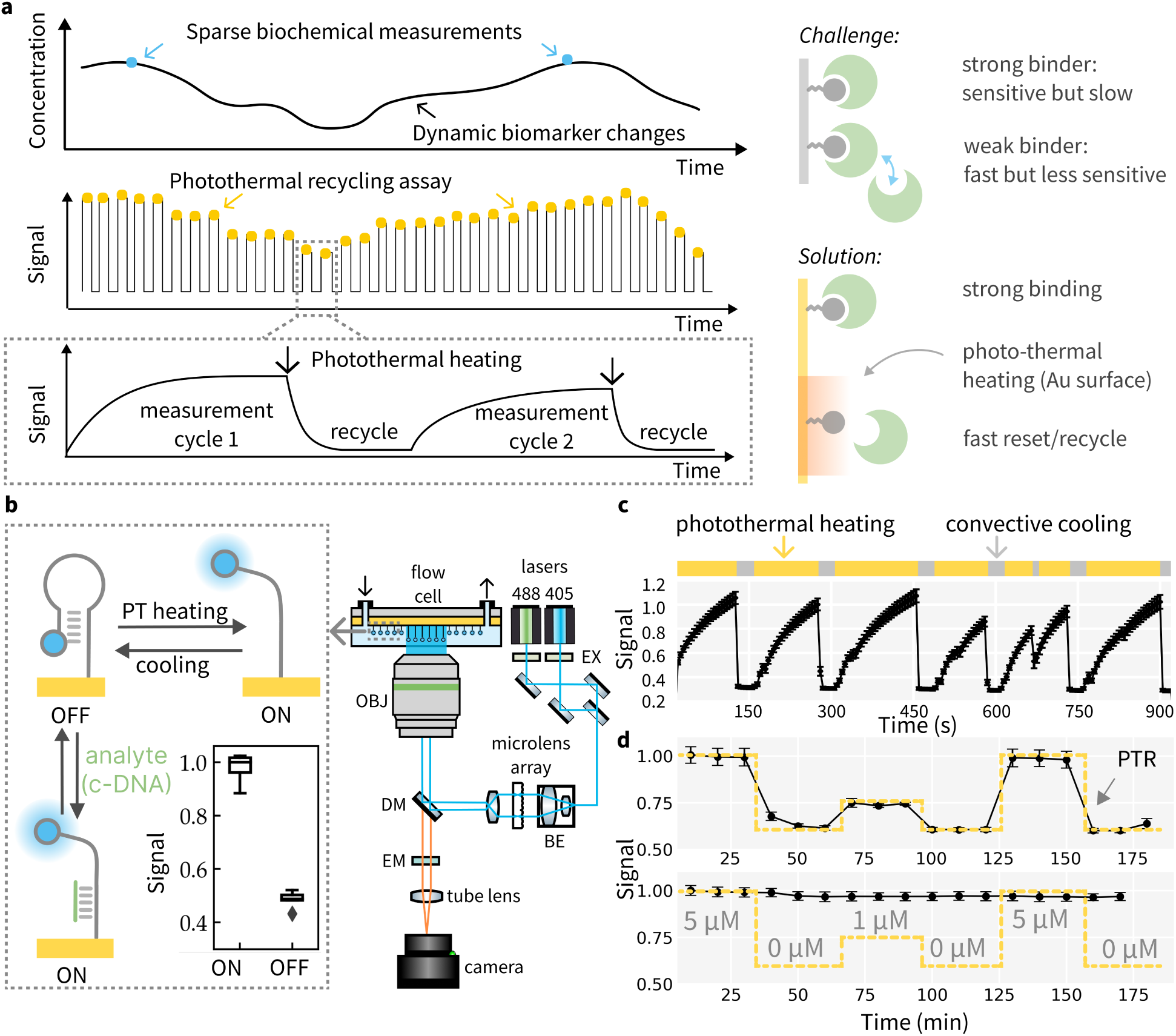
Photothermal recycling of bio-molecule probes. (a) PTR sensor working principle: localized, rapid heating disrupts tight binder-analyte interaction and enables frequent sampling of biomarker levels. (b) Hardware setup and a model DNA hairpin system to demonstrate the PTR assay: heating or complementary binding with c-DNA unfolds the hairpin structure that induces an enhanced fluorescence. Removal of the c-DNA strand or cooling resets the haripin.(EX: excitation filters, BE: beam expander, DM: dichroic mirror, EM: emission filters, OBJ: objective)(c) Repeated PTR cycles with the DNA hairpin show rapid cooling and steady fluorescence emission. (d) PTR sensor for oligonucleotide analyte: PTR enables continuous detection of the c-DNA target (top panel), while a low ionic strength solution (e.g. water) cannot recycle the sensor for repeated measurements (lower panel).

To achieve fast sensor temporal response without sacrificing sensitivity, disruptive force can be introduced to facilitate analyte dissociation and regenerate binding elements. Current biosensor regeneration techniques encompass a range of strategies, including surface re-engineering and re-functionalization, chemical treatments, allosteric modulation of binders through light or heat, and the application of electric or magnetic fields. [12–14] Methods that enable in situ regeneration of sensing interfaces are of particular interest for continuous biochemical monitoring, but many existing approaches face limitations in applicability for diverse sensor formats and clinical applications. Few strategies have been validated for regenerating sensing interfaces in continuous sensing applications. The temperature increase in bulk solution is unnecessarily slow and often requires specialized temperature control instrumentation. Localized heating techniques, which have shown efficacy in applications such as cancer photothermal therapy[15–18], offer a more practical alternative. To date, there has been no prior work that utilizes localized thermal energy as a recycling mechanism for continuous biochemical sensing applications.

In this work, we introduce a novel sensing mechanism, termed photothermal recycling (PTR) biosensing, to achieve highly sensitive and continuous biochemical quantification.(Fig.1a) Plasmonic materials, such as metallic nanoparticles and nanostructures[19], can absorb light irradiation and dissipate the energy as heat through non-radiative decay pathways[20], producing rapid and localized heating that modulates adjacent biomolecular processes and functions. Recent studies have demonstrated the application of photothermal heating in ultra-fast PCR[21–25] and cancer photothermal therapy[15–18]. In the PTR biosensing scheme, we propose to leverage photothermal heating as a disruptive force to facilitate the dissociation of tight analyte-binder complexes, thus recycling sensor elements for frequent measurements with minimal readout delays. This approach incorporates unique physical properties of plasmonic materials to modulate biomolecular recognition processes, which improves biosensor temporal response without sacrificing analytical sensitivity. Here, we report the design, optimization of PTR biosensing mechanism and demonstrate experimental results of continuous, ultrasensitive detection of diverse analytes such as proteins, oligonucleotides, and small molecules.

## 2. Results

### Photothermal Recycling of Biomolecular Probes

To confirm photothermal effect could reversibly modulate the biomolecular recognition process, we used DNA hairpin as a model system to experimentally demonstrate the effects of temperature cycling on bio-conjugation, photo-stability of fluorophore labels and biomolecular binding. Fig.1b shows the optical instrument setup for PTR sensor. DNA-based reagents are well known for their thermal stability to maintain structural integrity through multiple cycles of denaturation and annealing[26].

A DNA hairpin (Table.S1, a) is fluorescently labeled with Alexa488 dye on the 5’ end and covalently immobilized on the gold (Au) surface through a gold-sulfur bond[27, 28]. In this study, the gold-sulfur bond created using cyclic disulfide (Fig.S1) demonstrates comparable thermal stability to that of bonds formed using thiol or acyclic disulfide. Furthermore, cyclic disulfide exhibits greater chemical stability, particularly in the presence of competing thiols that are typically used for surface passivation[27].

Due to the strong plasmon quenching effect, the fluorophore label on DNA will demonstrate a distance-dependent intensity change at the Au surface[29, 30], which produces a fluorescence readout signal corresponding to the conformational states of DNA. In the “off” state, the DNA hairpin structure positions the fluorophore close to the Au surface, while in the “on” state the hairpin structure is disrupted and repositions the fluorophore away from the surface, leading to an increase in fluorescence signal. Using an invading short oligo to induce strand displacement, we confirmed that the model DNA system behavior is consistent with previously reported results, and we can observe a significant increase in fluorescence after hairpin opening. To ensure uniform heating across the field of view, we integrated a beam shaper into the excitation path, which converts the Gaussian illumination profile to a top-hat beam profile. (Fig.1b)

Au thin film absorbs light strongly at the plasmon resonance frequencies that lead to photothermal heating.[21] When illuminated continuously without convective cooling (no flow), the heating effect will induce a gradual fluorescence increase similar to using a displace-ment strand. This effect is due to thermal denaturation of the stem region of the DNA hairpin. When flow is introduced, convective cooling near the surface allows hairpin structure to reform which quenches fluorescence. Under mild illumination (see Methods for details), the photothermal heating and cooling process can be repeated multiple cycles without noticeable damage to the surface or the DNA beacon (Fig.1c). Although laser illumination could induce photo-bleaching of fluorophores[31], the observed cyclic fluorescence changes indicate that photothermal recycling is compatible with biomolecule-fluorophore construct for optical sensing applications. The surface and local temperature profiles resulting from localized photothermal heating and thermal transport were also measured using the temperature coefficient of resistance (TCR) method and modeled in COMSOL (Fig.S2). The temperature increase at surface was found to be around 22 °C, which is adequate to disrupt the DNA secondary structure.

To demonstrate the concept of photothermal recycling, we used an invading complementary DNA (c-DNA, 24 bp) strand as a mock analyte to simulate tight analyte-binder complex formation. (Fig.1b) The c-DNA strand exhibits a strong binding affinity and forms a stable complex with the DNA beacon, producing a sustained increase in signal. Even in low ionic strength conditions, such as deionized water, the displacement strand remains strongly hybridized to the DNA hairpin on the surface. However, by incorporating photothermal heating between measurement cycles, the DNA hairpin can be effectively recycled for binding. (Fig.1d) This approach allows accurate measurement of analyte concentrations over time and precise monitoring of their dynamic fluctuations.

### Digital Assay with Improved Sensitivity

Although the PTR DNA hairpin sensor demonstrates DNA concentration tracking, its dynamic range from µM to mM is inadequate for low-abundance analytes that typically have a physiological range in pM [32]. This limitation on analytical sensitivity is due to the high background of the fluorogenic assay format. To reduce assay background and enhance sensitivity, we adopted a digital assay format that quantifies discrete ON/OFF signals rather than continuous fluorescence changes. [33–39] (Fig.2a) Here, functionalized fluorospheres or beads will immobilize on surface via sandwich complexes in the presence of target molecules. The quantity of bound fluorospheres correlates with analyte concentrations. In theory, a digital bead assay utilizing single DNA sandwich complexes could achieve sensitivity to sub-fM analyte concentrations [40], given that sandwich formation in the absence of analyte is infrequent.

**Fig. 2:**
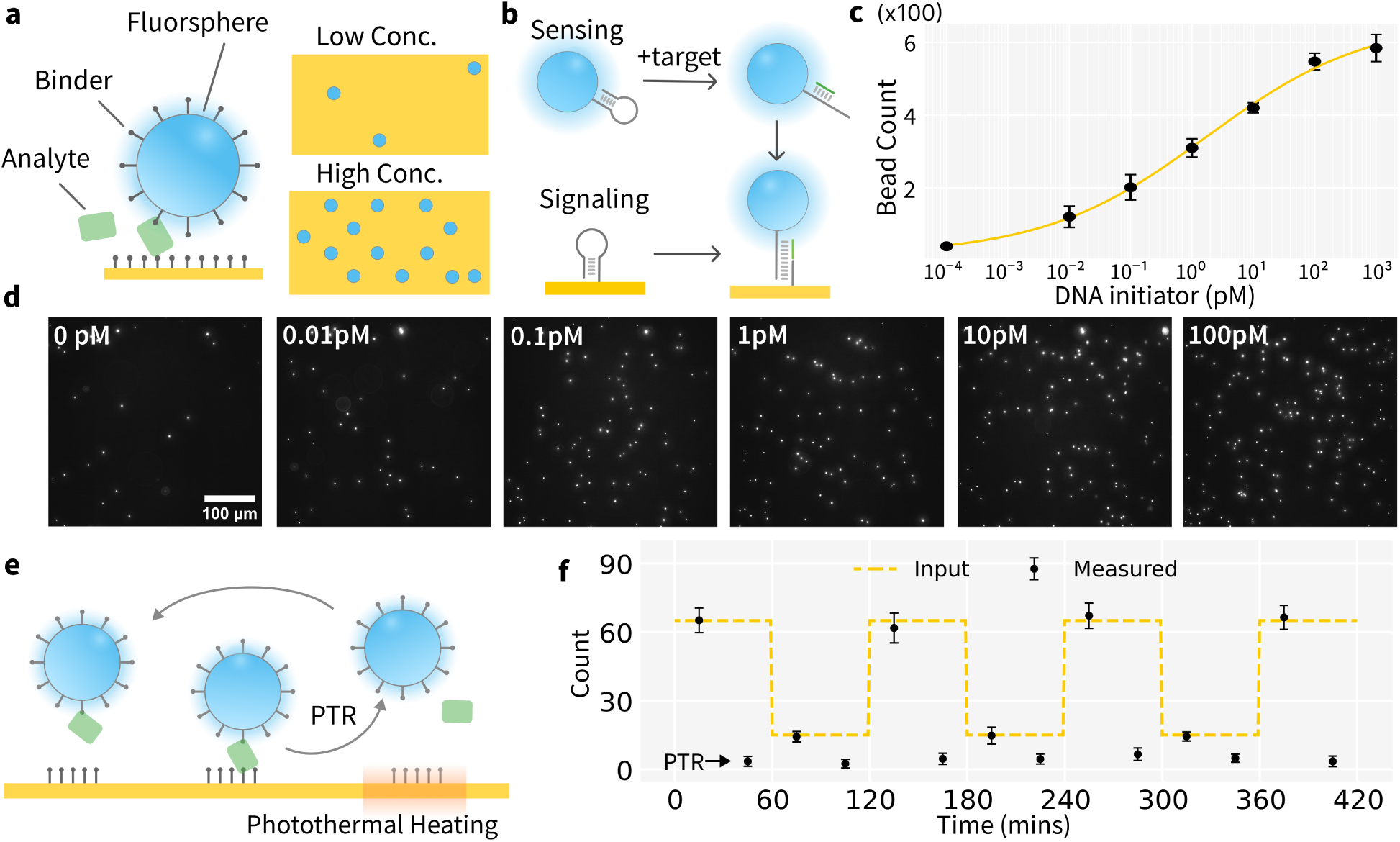
Digital bead assay provides ultra sensitive detection. (a) In digital bead assay, the analyte concentration is measured by counting the number of beads on the surface. (b) The assay mechanism for oligonucleotide analyte. Two DNA hairpins are initially self-hybridized, preventing bead-surface attachment. Target addition opens up one hairpin, which enables bead immobilization on the surface. (c-d) Binding curve and fluorescence images of the digital bead assay showing bead counts increasing as the DNA initiator concentration increases. (e-f) Using photothermal heating to recycle reagents and sensing surface enables repeated analyte measurements over multiple cycles.

We designed a generalizable digital assay mechanism as shown in Fig.2b. (Table.S1) A thiolated DNA hairpin (H1) is immobilized on surface, while a biotinylated complementary hairpin (H2) is attached to an avidin-coated fluorosphere. These two strands provide two essential functions: sensing the target analyte and producing a readout signal. In the absence of the initiator target, both hairpins (18bp stem) are topologically locked from hybridization[41], which prevents fluorospheres binding to the surface. When the initiator is present (24bp), it opens H2 via strand displacement, which then hybridizes with H1 (24bp stem), anchoring the fluorosphere to the surface (Fig.2b). This assay, based on the DNA strand displacement and hybridization chain reaction[41, 42], allows sub-pM DNA detection (Fig.2c and d, Fig.S3). Despite some background noise due to signal leakage[42–44] from hairpin, the PTR digital assay can detect target concentrations from low fM to high pM, spanning a dynamic range of about four orders of magnitude. (Fig.2c and d) The negative control with blank sample concentration is plotted as a pseudo low concentration in the binding curve. This increased sensitivity and dynamic range result from reduced background from non-specific binding compared to fluorogenic assays. Increasing the sampling area can also reduce noise from surface variability and passivation defects. As a result, digital bead assays complement the dynamic range achievable with continuous flow assays[9], which detect concentrations from low pM to high nM.

### Photothermal Recycling of Digital Assays

Although digital assays are ultrasensitive, they cannot respond to dynamic changes in analyte concentration. Combining PTR with digital assay format enables ultrasensitive, continuous analyte tracking. (Fig.2e) We functionalized fluorospheres with complementary DNA (Table S1, g&h) to mimic bead attachment to the surface in the presence of analyte. Photothermal heating denatures the DNA duplex, and buffer advection in the flow cell removes the detached beads. This process recycles the sensing surface for repeated measurements.The bead count after each PTR reset step can serve as an internal quality control of the surface in different matrices. (Fig.2f; Fig.S4) Video.S1 demonstrates the dynamic bead count changes within a measurement cycle on the Au surface while Fig.S5 plots the bead count over time. Effective bead detachment was observed with the 405 nm and 488 nm lasers at full illumination for 2mins, while only minimal bead detachment occurred under 640 nm full illumination (Fig.S6-7). It confirms the lifting is due to the photothermal effect. At moderate illumination, no lifting is observed with either 405nm, 488nm or 640nm laser. We chose 405nm at full illumination for photothermal heating and 488nm at moderate illumination for imaging.

We studied the relative strength of binding and photothermal disruption between beads and the Au surface to optimize specific binding while reducing non-specific binding. The DNA-bead system relies on duplex hybridization for signaling, making the strength of specific bead-surface interaction highly tunable through adjusting DNA sequence (i.e., affinity) and density (i.e., avidity). It is suitable for probing effects of binding affinity and avidity, and optimizing photothermal heating conditions that could be generalized for both DNA-based reagents (e.g., aptamers) or binder scaffolds. (Fig.3a) The disruptive force from photothermal heating depends on the total input energy (laser power integrated over time), absorption efficiency, and photothermal conversion efficiency. It is also affected by the thermal transport at the Au-liquid interface. We measured and modeled the temperature increase at this interface using multiphysics simulation, as shown in Fig.S2.

**Fig. 3:**
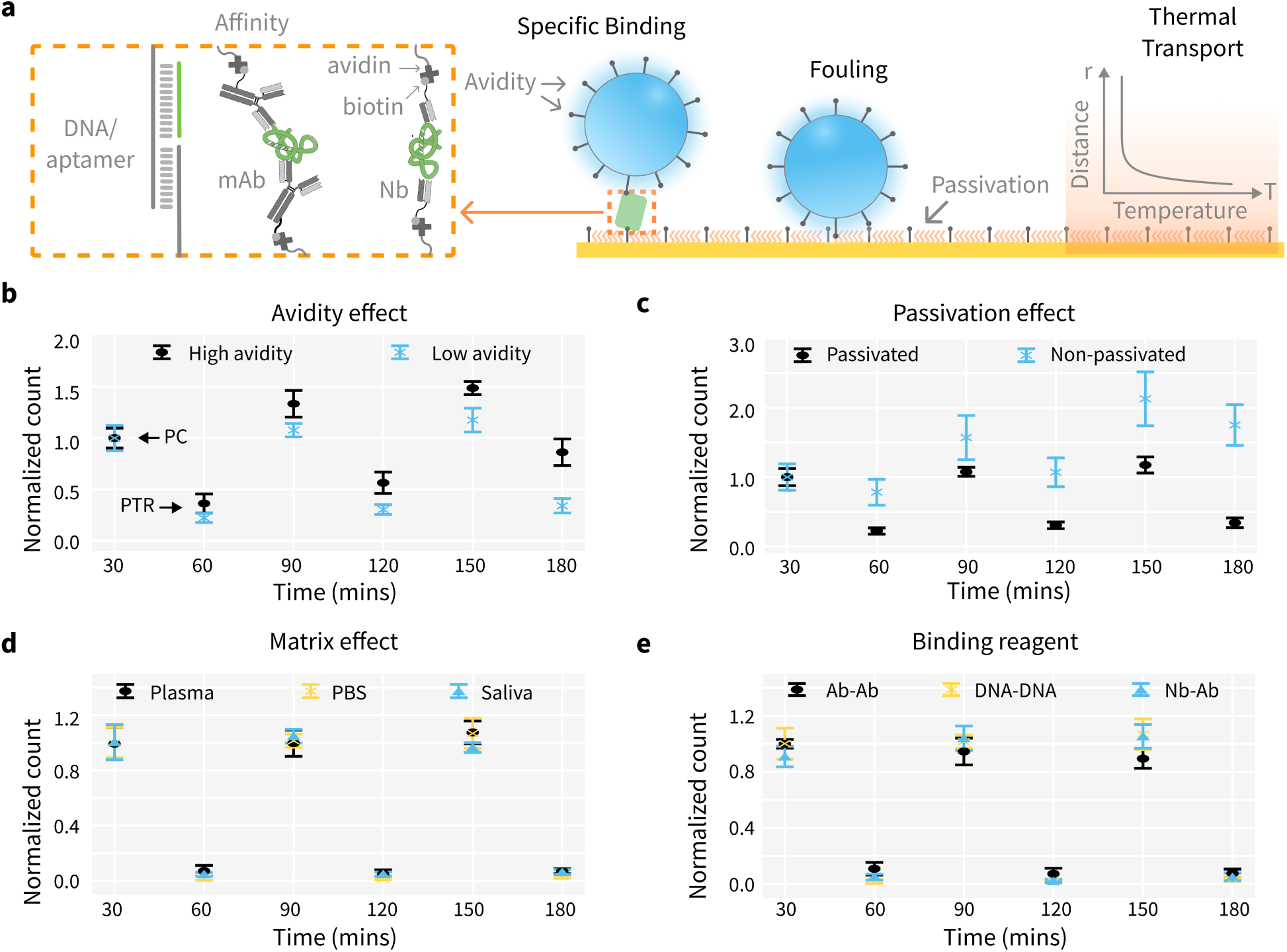
Digital PTR Assay Optimization. (a) Key factors affecting analytical performance, assay longevity and recycling efficiency are optimized to improve specific binding and minimize fouling. mAb: monocloncal antibody; Nb: nanobody. (b) Avidity effect: DNA bead with high avidity binds tighter and requires a stronger external force to remove. The positive controls (PC) with DNA-modified beads are run for three cycles. In each cycle, a measurement is followed by a photothermal recycle (PTR). (c) Passivation effect: DNA beads will accumulate on a non-passivated surface even after heating cycles, while a passivated surface remains clean. (d) Matrix effect: Photo-thermal recycling works in multiple matrices including plasma, PBS and saliva. (e) Binder: Various binding scaffolds have been evaluated to be compatible with PTR.

We demonstrate that, for fixed disruptive force and avidity, decreasing the affinity by lowering the buffer ionic strength enhances photothermal lifting efficiency (Fig.S8). For fixed disruptive force and affinity, lower avidity also leads to higher photothermal lifting efficiency (Fig.3b). The positive controls (PC) are ran for three cycles. In each cycle a measurement is followed by a recycle (PTR). The surface can be effectively recycled without obvious accumulation with lower avidity case. Surface passivation significantly enhances photothermal lifting efficiency, as non-passivated surfaces tend to retain fluorospheres, hindering their detachment. Using 6-mercapto-1-hexanol (MCH) for passivation in PTR assays prevents bead accumulation, whereas severe accumulation occurs on unpassivated surfaces even after heating.(Fig.3c) Under optimized conditions, photothermal recycling can be effectively applied in various matrices without significant accumulation of fluorospheres on the Au surface. The PTR bead assay has been demonstrated in plasma, PBS, and saliva without observable accumulation. (Fig.3d) Various binding reagents have been evaluated within the PTR scheme, including DNA-DNA interactions, antibody-antigen interactions, and protein-nanobody interactions. Results demonstrate that the sensor surface can be effectively regenerated for continuous detection for these reagents. This supports the potential for further development of PTR sensors utilizing protein binders. (Fig.3e)

### PTR Digital Assay for Protein and Small Molecule Detection

For the detection of protein and small molecule targets using PTR digital assay format, we designed two generally applicable mechanisms using either oligonucleotide or a proteinbased binder. For protein targets that usually have multiple epitopes, we adopted the standard sandwich format using two paired binders on bead and surface. We use thrombin and thrombin-binding aptamers[45] (TBA, Table S1 e&f) to demonstrate the sandwich PTR assay. Thiolated aptamer TBA1 is covalently immobilized to Au surface and biotinylated TBA2 is attached to an avidin-coated fluorosphere. Sandwich complex formation anchors the fluorosphere to the Au surface for digital assay.(Fig.4a) This assay format enables sub-pM thrombin detection in both buffer (Fig.S9) and diluted serum (Fig.4b). When dynamically changing the spike-in thrombin concentrations (input), the measured results exhibited excellent alignment with input concentrations, confirming the PTR sensing design can preserve digital assay sensitivity while enabling continuous measurement.(Fig.4c) The standard curve is plotted as log-log to emphasize the linear relation in the assay range.

**Fig. 4:**
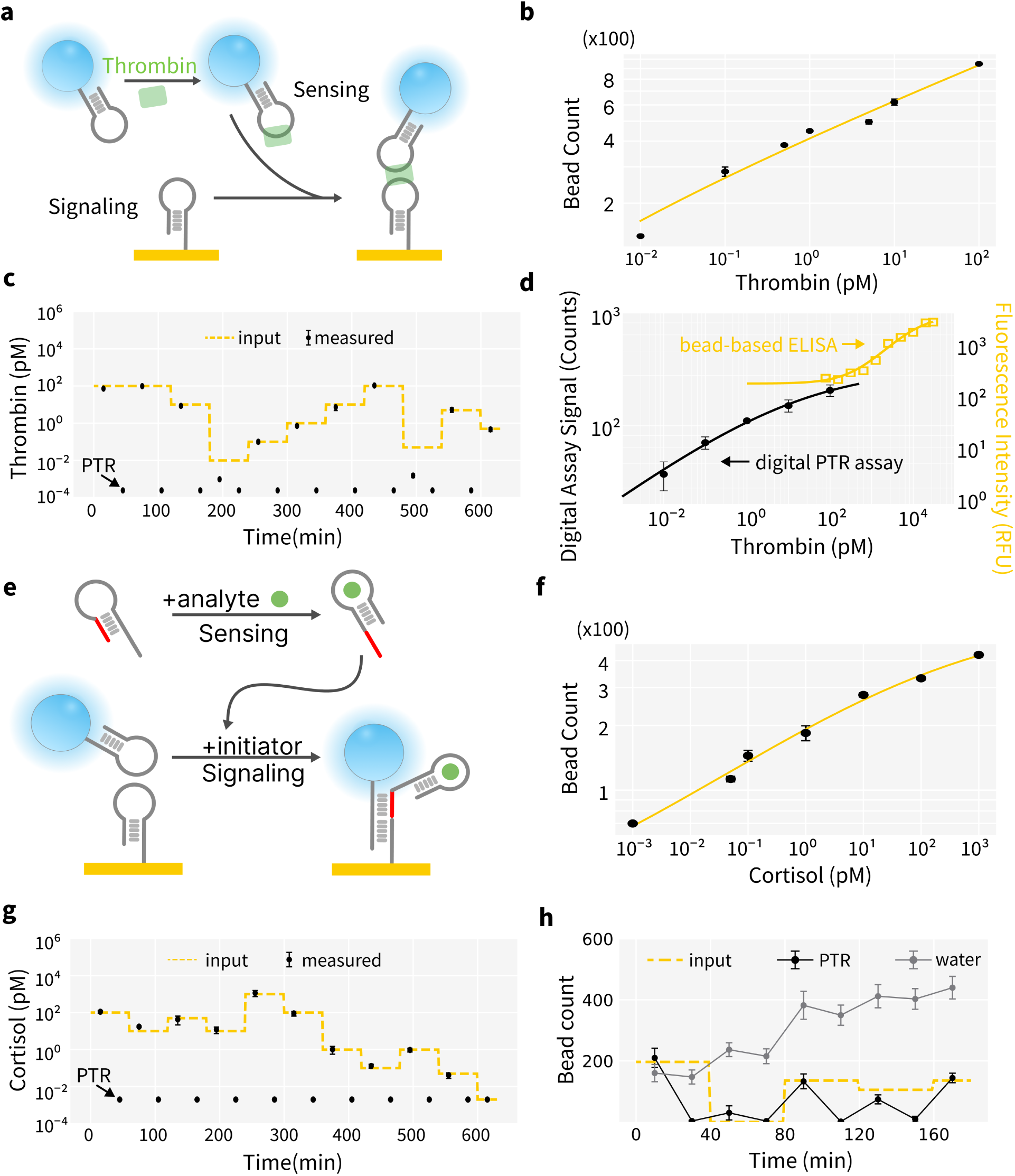
PTR Enable Continuous Ultrasensitive Detection of Biomolecules. (a) Sandwich PTR assay format. (b-c) Ultrasensitive continuous detection of thrombin in diluted serum. (d) Comparison of PTR digital assay versus continuous bead-based ELISA assay[9]. (e) PTR assay format for detecting small molecules. (f-g) Ultrasensitive continuous detection of cortisol in saliva. (h) PTR effectively recycles the surface for continuous measurements while washing with DI water alone cannot reset the surface for repeated measurements.

For small molecule targets such as metabolites or small peptides, it’s challenging to find two epitopes for sandwich assay format. We combined our digital assay mechanism(Fig. 2b) with advances in DNA hybridization chain reaction[46, 47] and structure-switching aptamer design[48] to develop small molecule PTR assay. We use cortisol as a model analyte to demonstrate assay performance. (Table.S1 j&k) Without cortisol, the cortisol aptamer-initiator strand adopts a hairpin conformation, sequestering the initiator sequence and preventing signal generation. In the presence of cortisol, conformational switch of a cortisol aptamer exposes the initiator sequence that enables H1 and H2 to hybridize and anchor fluorophores to the surface. (Fig.4e) This study demonstrates continuous, ultrasensitive monitoring of cortisol in buffer (Fig.S10) and artificial saliva (Fig.4f&g), detecting concentrations from low fM to high pM, spanning four orders of magnitude. A log-log plot is used to emphasize the linear relationship within the assay range. As a control to demonstrate the necessity of photothermal disruption, we compared using DI water and photothermal heating to recycle the digital assay surface. As shown in Fig.4h, using only DI water the digital assay surface is gradually saturated while the PTR sensor surface retains it’s ability for analyte quantification.

### PTR Biosensing for Inline Extracellular ATP Monitoring

We demonstrate the utility of PTR biosensing for inline biochemical monitoring of biological processes. We use Adenosine triphosphase (ATP) in bacterial culture as a model system as it’s an important metabolism marker and is released extracellularly via metabolic activity, stress responses, membrane disruptions or cell lysis.[49] In E. coli cultures, extracellular ATP (eATP) peaks at the end of the log phase and declines in the stationary phase due to uptake or degradation at the cell surface [50], as bacteria can actively internalize and utilize extracellular ATP.[51] To monitor ATP levels during cell culture, we developed an inline assay setup that can frequently sample cell culture for measurements. (Fig.5a) We adapted the DNA digital bead assay for ATP detection by extending the DNA initiator into an ATP-binding aptamer switch (ATP initiator, Table.S1,I), similar to previously reported ATP aptamer switches [42]. DNA hairpins (H1, Table.S1,c; H2, Table.S1,d) were displayed on both the Au surface and fluorospheres the same way as in PTR digital assay mechanism(Fig.4e). Upon ATP binding, the initiator region is exposed, triggering a DNA strand displacement between H2 and H1, and enabling fluorosphere binding to the gold surface. (Fig.5b) The thermodynamic and kinetic properties of aptamer switches can be engineered to meet the dynamic range requirement of inline biosensing[52]. Assay binding curve and spike-in experiment results show the PTR assay can achieve continuous ultra-sensitive ATP detection in buffer(Fig.5c).

**Fig. 5:**
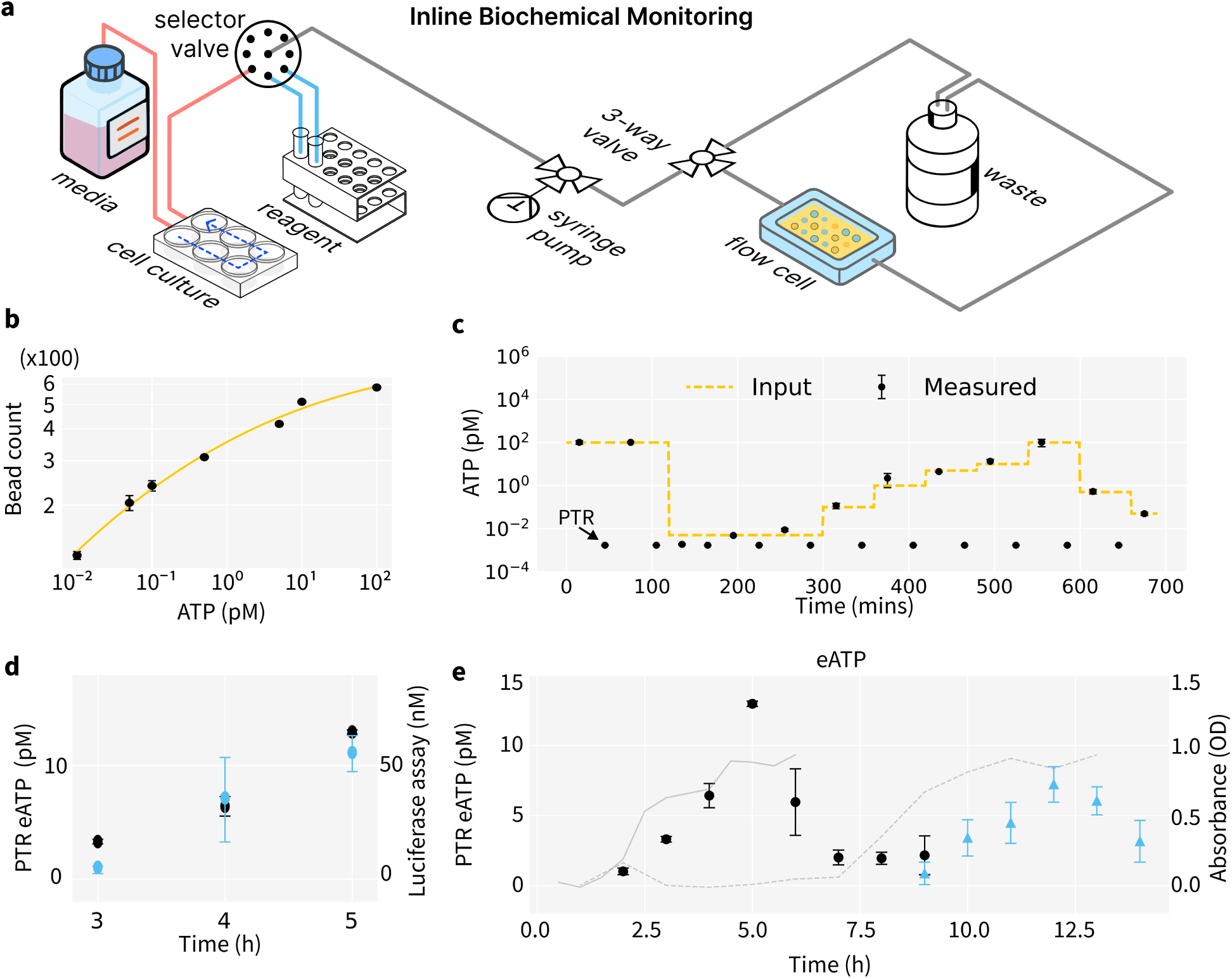
Application of PTR Biosensors for Inline Biochemical Monitoring. (a)Measurement Setup for extracellular ATP in cell culture. (b) Binding curve of ATP detection using the inline digital PTR assay. (c) Continuous detection of ATP demonstrated in buffer (d)The PTR eATP measurement (black) is cross-validated with the ATP luciferase assay (blue). The original sample is diluted 5,000 times for PTR measurement. (e) Monitoring dynamic change of eATP in E.coli culture. The trend of eATP (scatter) is plotted with optical density measurement (grey line) over cell culture stages. The trend of eATP in E.coli culture media without pre-stabilization (blue scatter) shows a delayed onset compared to the pre-stabilized group (black scatter).

We further cross-validated our PTR assay with a commercial luciferase ATP assay (Fig.5d), which shows good agreement between our PTR assay with the commercial assay. Our results show similar to previous studies[50, 51], extracellular ATP levels reach their maximum towards the end of the logarithmic growth phase and decline as the cells enter the stationary phase (Fig.5e, black scatter). A decrease in extracellular ATP concentration is observed, which indicates that the DH5a E.coli strain can uptake eATP from culture media. The measured ATP trend aligns with that measured with the ATP luciferase assay (Fig.S11). Furthermore, a delayed onset of the eATP trend is observed in a group of E.coli without pre-stabilization after thawing from frozen stock (blue scatter).

These results indicate that the eATP measurement can profile the proliferative status to culture condition changes. The PTR sensor demonstrates high sensitivity and a lower detection limit, requiring only a minimal sample volume in a few micro-liters per assay. This offers several advantages. First, small sample volume does not interfere with the cell culture process, eliminating the need for additional feeding procedures. Second, diluting the sample with buffer can enhance the device’s longevity, as nonspecific binding and the accumulation of fluorospheres in complex media conditions are often more severe than in buffer which can lead to passivation failure. These combined benefits simplify and improve the continuous monitoring of extracellular ATP (eATP) in bacterial culture media on the PTR sensor. Cultures can be initiated with a smaller volume, allowing for multiple measurements using the same device.

## 3. Discussion

Current biosensors face limitations in continuous detection of analytes below pM concentrations in biofluid matrix due to slow diffusion and binding kinetics of the affinity-based immunoassay format[6, 53–55]. Aptamer-based electrochemical sensors are often used for continuous monitoring applications to improve readout frequency and speed, but the relatively low-affinity aptamer-analyte interactions limit the sensitivity of ensemble-averaged electrochemical readouts. [56–59] On the other hand, digital bead assays offer exceptional sensitivity[39], but their adaptation for continuous detection has been limited. Existing continuous digital assays[53, 54], such as those leveraging Brownian motion of tethered particles, face similar challenges to balance assay speed and sensitivity.[55]

Our sensing mechanism design circumvents the sensitivity/speed limitation of biomolecular binding/dissociation by leveraging unique photo- and thermo-physical properties of plasmonic materials to modulate the spatial and temporal characteristics of the molecular binding process. Plasmonic materials that can concentrate and convert light energy through light-matter interactions offer a unique capability to control the energy transfer and mass transport process during biomolecular interaction.[60, 61] Our results show that a functionalized Au surface under moderate illumination power could achieve rapid, highly localized temperature increase of approximately 21 K near the bio-nano interface surface. This local temperature gradient is sufficient to disrupt molecular interactions, and subsequent convective cooling could effectively recycle the binders and sensing surface for the next detection cycle.

Our multi-hour PTR assay confirms DNA-based reagents, which is known to have exceptional thermal stability[26], can retain their structural integrity through repeated cycles of denaturation and folding. This enables advances in DNA technology, such as rational design of novel binders, switches and leak-resistant signaling mechanisms with programmable binding energy and thermodynamic properties [62–66], to be integrated into the PTR sensing mechanism. At the same time, Au-DNA surface is commonly in the electrodes of aptamer electrochemical sensors, which allows the large body of surface passivation studies[67–71] for implantable electrochemical sensors to be leveraged in the development of the PTR sensing scheme. In addition to DNA-based reagents, our results also show protein-based reagents such as antibody and nanobody (Fig.4e) is compatible with thermal cycling. Recent developments in the computational design of thermally stable protein binders[72–74] offer enhanced resilience under heating. Integrating these advanced protein binders with PTR digital biosensors could broaden detection capabilities to a wider variety of target molecules.

## 4. Conclusions

This study presents a novel sensing mechanism that integrates photothermal recycling with digital bead assays to enable ultrasensitive, continuous detection of low-abundance biomarkers in the fM to pM range, including nucleic acids, proteins, and small molecules. Our results show that localized temperature gradients could effectively modulate the kinetics of molecular binding over multiple heating/cooling cycles for both nucleic acids and protein binders. We demonstrate the utility of this sensing approach in the inline monitoring of metabolites in a cell culture. This biosensing mechanism is highly versatile and has the potential to be integrated with various optical sensing platforms for applications in bioprocess monitoring, biomedical diagnostics, and therapeutics. Translation to these applications requires further research to systematically evaluate the surface chemistry and photothermophysics of different materials under diverse experimental conditions (e.g., body temperature) over an extended period, and assay validation with application-specific matrices (e.g., blood, culture media) and binding reagents.

## Acknowledgment

The authors acknowledge financial support from the National Institute of Biomedical Imaging and Bioengineering Award R21EB031455, National Science Foundation Biosensing Program Award 2339756, and University of Florida Research Opportunity Seed Grant.

## Author Contributions

Y. T., Y. L., W. W., M. C. performed all assay design and experimental characterization. Y. L., Z. Q., Q. Z. performed surface temperature measurement and modeling experiments. Y. T., Y. L., Z. Q., J. P. contributed to the initial paper draft. Y. T., J. N., C. D., and J. P. consolidated all the writings, reviewed and revised the final manuscript. J. P. conceptualized the research topic and supervised the research and writing. All authors reviewed the final version and agreed on its submission.

## Additional information

Correspondence and requests for materials should be addressed to Jing Pan.

## 5. Methods

### Materials

Glass slides (25×75×1.0mm, Precleaned, Premium Plain Microscope Slides, Cat. No. 125444, Fisherbrand), cover slips (24×60mm No.1.5 thickness, epredia), PBS (PhosphateBuffered Saline, without calcium and magnesium, 1x, Corning), and Tween-20(BP337-100) were obtained from Fisher Scientific. Double-sided adhesive tapes were processed using a desktop cutter to obtain the pattern of the flow channel. The ssDNA oligonucleotides (standard desalting and HPLC purification for chemically modified DNA) were purchased from IDT (Integrated DNA Technologies). The sequence and modifications of all ssDNAs are listed in Table S1. Tris(2-carboxyethyl)phosphine hydrochloride Solution (TCEP, 0.5M, 646547-10X1ML), MCH (6-Mercapto-1-hexanol, 451088-5ML), FBS (Fetal bovine serum, heat inactivated), and artificial saliva(1700-0305) were obtained from Millipore Sigma. The Avidin modified flurospheres (Molecular Probes FluoSpheres NeutrAvidin-Labeled Microspheres, 1.0 *µ*m, yellow-green fluorescent), human *α*Thrombin Native Protein (RP43100,) and ATP solution (100mM, FERR1441) were obtained from Thermo Fisher. The cortisol standard solution is obtained from a cortisol parameter assay kit (KGE008B, Biotechne). The library efficiency DH5*α* competent cells are purchased from Thermofisher and a pUC19 DNA plasmid is encapsulated for hepcidin production. BacTiter-Glo™ Microbial Cell Viability Assay kit (G8230) is obtained from Promega.

### Hardware Setup

The imaging system comprises a TIRF Nikon microscope integrated with a custom-built laser launch system (Fig.1b). This system employs two laser lines: 405 nm (30mW) for photothermal heating and 488 nm (6 mW) for imaging. A Top-Hat beam shaper is incorporated to provide uniform illumination over a square region, with precise details of the illumination area and laser power specified for optimal performance. The lasers are initially expanded with a beam expander to the size required by the TOPAG beam shaper. The light is then uniformly redistributed into a square profile and relayed into the sample plane using a tube lens and objective.

### Surface preparation

The inlet and outlet holes are drilled with 1mm round ball head diamond grinding bits on standard microscope slides (L=75mm W=25mm H=1mm) prior to gold coating. Standard glass microscope slides are cleaned using plasma and piranha solution prior to use. A thin 5-10 nm layer of chromium/nickel is then sputtered onto the cleaned slides to enhance adhesion, followed by the deposition of a 100 nm layer of gold through sputtering. Sputteringwas done using a sputter deposition machine (KJL CMS-18 multi-source).

The flow channel is assembled by sticking a cover slip to the gold-coated slides using the patterned adhesive. The holes are drilled into the PDMS connectors to accommodate the metal pins and connect to the tubing. These PDMS connectors are then affixed to the inlets and outlets, preparing the channel for use.

For DNA surface preparation, first, the thiolated DNA oligonucleotide is reduced prior to use with TCEP. 4 *µ*L of 100*µ*M thiolated DNA is mixed with 4 *µ*L 500*µ*M TCEP for 1.5 hour. Then, the cleaned gold substrate is immersed in a 1.0 µM solution of thiolated DNA oligonucleotide in 1*×* PBS buffer (pH 7.0) for a specified duration, ranging from 2 hours to overnight. Following incubation, rinse the substrate with 1*×* PBS buffer. In thrombin assays, the thrombin aptamer TBA1 (Table S1.e) is immobilized on Au surface. In DNA initiator, ATP and cortisol assays, H1 (Table S1.c) is immobilized on Au surface.

The surface is then passivated with 6-mercapto-1-hexanol (MCH). The channel is immersed in a 10 mM solution of MCH in 1*×* PBS at room temperature for 2 hours to passivate the surface. Afterward, perform a final rinse with 1*×* PBS buffer. The sensor is then ready for use.

Previous studies report a surface coverage of 5.2 (*±*0.8) *×* 10^12^ molecules/cm^2^ for probes immobilized in an HS-ssDNA/MCH mixed monolayer, as measured by SPR under similar preparation conditions[7], and 5.7 (*±*0.05) *×* 10^12^ molecules/cm^2^ as determined by ^32^Pradio-labeling experiments[75].

### Fluorspheres labeling

Incubate 4 *µ*L of 1 mM biotinylated DNA reagent with 8 *µ*L of 1% fluorsphere (FS) solution in 100 *µ*L of PBS for 15 minutes. Next, add 900 *µ*L of 1*×* PBS and 1% Tween 20 to the mixture, and mix thoroughly by vortexing. Centrifuge the mixture at 5,000 rpm for 30 minutes, then carefully discard the supernatant. The beads are redissolved in 1*×* PBS buffer, and the beads are ready for use. In thrombin assay, thrombin aptamer TBA2 (Table S1.f) is labeled on FS. In DNA initiator, ATP and cortisol assays, H2 (Table S1.d) is labeled on FS.

### PTR digital assay

The DNA initiator, human *α*thrombin, cortisol and ATP are prepared in 1*×* PBS buffer with a 10-fold serial dilutions from 10fM to 1nM. The human *α*thrombin is also prepared in 10% fetal bovine serum (diluted with 1*×* PBS buffer) with a 10-fold serial dilutions from 10fM to 1nM. The cortisol solution is also prepared in artificial saliva with a 10-fold serial dilutions from 10fM to 1nM.

For a standard sandwich photothermal recycling (PTR) digital assay cycle, begin by flushing the surface with buffer. The flow rate is set at 2000 *µ*L/hr for 200ul buffer. Incubate 100ul of conjugated fluospheres from previous step with 100ul solution containing different concentrations of target molecules for 15 mins by gentle vortexing and mixing. In ATP and cortisol assays, 4 *µ*L of 1 mM ATP initiator and cortisol aptamer complex are spiked in respectively. Introduce the prepared beads and incubate for another 15 minutes at a flow rate of 1000 *µ*L/hr, followed by an additional buffer flush at a flow rate of 2000 *µ*L/hr. The surface is then ready for imaging to acquire signals. Multiple locations are scanned sequentially on Au surface and the positions are recorded. For photothermal recycling, flush the surface with water at a flow rate of 2000 *µ*L/hr. Expose the surface to a 405 nm laser with 14000mW/cm^2^ for 2 mins while maintaining a continuous flow of deionized water at 1000 *µ*L/hr. The same locations are regenerated and imaged sequentially to demonstrate the effect of PTR.(Fig.S12) The surface is then prepared for the next measurement cycle. Samples of different concentrations are measured using the PTR sensor, and the collected bead counts are correlated with the input concentrations using a four-point logistic model to generate the binding curve.

### PTR assay optimization

For optimization PTR assay, direct hybridization of DNA1rc (Table S1.h) labeled fluorspheres onto the DNA1 (Table S1.g) immobilized Au surface are used as a modal system. For avidity effect study, mix 0.1 *µ*L of 100 *µ*M (low avidity) or 10 *µ*L of 100 *µ*M (high avidity) biotinylated DNA reagent with 8 *µ*L of 1% fluorsphere (FS) solution in 100 *µ*L of PBS buffer solution for 15 minutes. The PBS buffer is diluted with diH2O to make 0, 0.01, 0.03, 0.05, 0.1, 0.3, 0.5 and 1*×* PBS buffer solution. Next, add 900 *µ*L of PBS buffer solution and 1% Tween 20 to the mixture, and mix thoroughly by vortexing. Centrifuge the mixture at 5,000 rpm for 30 minutes, then carefully discard the supernatant. The beads are redissolved in PBS buffer solution, and the beads are ready for flow into the sensor channel and test for lifting efficiency.(Fig.3b)

For passivation effect test, the PTR lifting effect is studied on Au surface with MCH passivation and without passivation.(Fig.3c) For matrix effect study, the DNA1rc fluorspheres are prepared in plasma, saliva and PBS buffer and tested for 3 cycles on Au surface.(Fig.3d) To study PTR regeneration effect regards the laser illumination time, 1 mins, 2 mins and 3 mins of PTR time are tested for lifting efficiency for 3 cycles.(Fig.S6) For PTR lifting efficiency regarding the laser wavelength, 2 mins of illumination of 405nm, 488nm and 640nm laser with 14000mW/cm^2^ on surface for photothermal heating is tested (Fig.S7).

For binding reagent test, antibody-secondary antibody, nanobody-protein and DNA-DNA interactions are studied. For immobilization of half antibody or nanobody onto Au surface, dissolve protein powder (1mg) in 100 mM PBS with 10 mM EDTA, pH 6.0 - 7.4. Mix the protein solution with TCEP so that the final protein concentrations of reduction mixtures is 10*µ*g/ml to 5mg/mL, TCEP concentration is in the range of 5mM to 125mM. Incubate the reduction mixture for 90 min at 37C and immediately flow into the sensor channel. For protein labeled fluorspheres, 4*µ*l 0.5mg/ml biotinylated secondary antibody or protein targets are mixed with 8*µ*l of 1% fluorsphere (FS) solution in 100 *µ*L of PBS buffer solution for 15 minutes. Next, add 900 *µ*L of 1*×* PBS. The mixture is directly used for PTR testing.(Fig.3e)

### PTR fluorescent assay

For the PTR fluorescent cycling experiment, after immobilization of the A488 labeled DNA beacon, the surface is flushed with water. The flow rate is set at 2000 *µ*L/hr for 200*µ*l water. The positive control is obtained by flushing the surface with 1N NaOH. For subsequent cycling, 5mW 488nm laser is for both imaging and photothermal heating and kept on for all time. The exposure time for imaging is set at 40ms. The flowrate of water for convective cooling is set at 200 *µ*L/hr. (Fig.1c)

For the PTR fluorescent assay monitoring DNA target, the positive control is also obtained by flushing the surface with 1N NaOH. 1 *µ*M and 5 *µ*M target DNA strand is prepared in the buffer with 1*×* PBS and 2mM Mg. In each measurement, the sample is flowed in and incubate for 10 mins. (Fig.1d)

### eATP collection and measurement

For DH5*α* competent cells were cultured overnight in LB broth at 37°C with shaking at 225 rpm. The overnight cultures were then diluted 1:100 in fresh LB broth and incubated at 37°C with continuous shaking. Aliquots were collected every 30 minutes during incubation, and the optical density at 600 nm (OD600) was measured to monitor growth. Following incubation, the bacterial cultures were centrifuged at 16,100 g for 5 minutes, and the supernatant was transferred to fresh tubes, which were stored at -80°C for later analysis. In the control experiment, ATP levels in the bacterial supernatant were assessed using the BacTiter-Glo Microbial Cell Viability Assay Reagent (Promega, Madison, WI). The collected cell supernatant is diluted (5,000 to 100, 000 times) based on the concentration range to be measured with the PTR sensor.(Fig.5d-e) An inline monitoring system is setup where the culture media is mixed with reagents inline and directly assayed in the PTR sensor chip.

### Photothermal Heating and Electrical Temperature Sensing

To experimentally validate the photothermal effect, we fabricated a microfluidic device consisting of a PDMS flow channel (35*×*15*×*0.15 mm^3^) sandwiched between two standard microscope slides with different lateral dimensions (35 *×* 32 *×* 2.5 mm^3^ and 35 *×* 15 *×* 2.5 mm^3^, Square Optics), as shown in Figure S2(a) and (b). A rectangular chamber (25 *×* 2.5 mm^2^) was carved from the center of the PDMS layer to contain the PBS solution. A Au (100 nm)/Cr (10 nm) bilayer was deposited by sputtering onto the glass slide surface, patterned with a central narrow strip (1 *×* 0.2 mm^2^) and four outer electrode pads for four-point probe measurement. This metallic pattern served as both the photothermal absorber and temperature sensor, with an initial room temperature resistance of 6.03 Ω. Prior to photothermal experiments, the temperature coefficient of resistance (TCR) of the metal strip was calibrated using a precision heating stage (Zhengketan KT-Z4019M4TR). The resistance was measured via four-point probe configuration by applying a small AC current (0.5 mA at 10 Hz, Keithley 6221) while monitoring the voltage with a lock-in amplifier (Stanford Research Systems, SR830). Temperature-dependent resistance measurements were collected over the range of 27-47*^◦^*C and analyzed by linear regression to determine the TCR.

Following calibration, the sensor surface underwent functionalization. First, 5 *µ*L of TCEP was dissolved in 1 mL of 1*×* PBS, and this solution was introduced into the chamber for 2-hour incubation at room temperature. After thorough rinsing with 1*×* PBS to remove residual reagents, a 20 mM MCH solution was prepared by mixing 897 *µ*L of 1*×* PBS, 100 *µ*L of ethanol, and 2.64 mg of MCH. Subsequently, 75 *µ*L of the MCH solution was injected into the chamber and incubated for an additional 2 hours at room temperature.

For photothermal heating experiments, a 405 nm LED (3546 mW total power, measured by power meter) illuminated the functionalized Au strip immersed in PBS solution, as illustrated in Figure S2(a) and (c). The Gaussian beam profile had an effective 1*/e*^2^ radius of 2.88 mm at the sensor surface (measured by Cinogy CMOS-1204 beam profiler via 2D Gaussian fitting), yielding an average power density of 13,608 mW/cm^2^. Temperature rise was monitored by applying a small probe current (0.1-0.5 mA, Keithley 6221) and converting the measured resistance change using the calibrated TCR. To correlate experimental results with uniform heating conditions, a scaling factor was determined through COMSOL simulations comparing Gaussian and top-hat beam profiles.

### COMSOL Simulation Details

A finite element model was constructed in COMSOL Multiphysics comprising two glass domains (10 *×* 10 *×* 2 mm^3^), a solution domain (10 *×* 10 *×* 0.15 mm^3^), and a gold sensor domain matching the experimental four-point probe geometry. The model was discretized into 523,345 elements, with mesh independence verified to ensure solution accuracy. Material properties were determined as follows: glass transmissivity at 405 nm was experimentally measured as 0.95 using a Thorlabs S121C power meter; PBS transmissivity was taken from [76]; gold absorptivity was obtained from [77]; and PBS thermal conductivity was sourced from [78].

Boundary conditions were implemented with all vertical glass surfaces fixed at ambient temperature to account for the large sample size, while thermal insulation conditions were applied to remaining boundaries. Heat generation was modeled by applying either Gaussian or uniform power distributions to the gold sensor surface. The simulated temperature distributions are presented in Figure S2(e) and (f). Volume-averaged temperature rise of the gold sensor was calculated for both beam profiles to determine the scaling parameter *δ* between Gaussian (experimental) and uniform (top-hat) illumination conditions. This scaling factor enabled conversion of experimental measurements to equivalent uniform heating conditions for comparison with theoretical predictions.

