## Supplementray Information for "Photothermal Recycling Biosensing for Continuous, Sensitive Molecular Quantification"

**Table S1. DNA Sequences**

| | Name | Sequence |
|---|---|---|
| a | DNA beacon | /5Alex488N/ TGC GAG AAA AAA AAA AAA AAA CTC GC /3ThioMC3-D/ |
| b | DNA initiator | CCC AGG TAA CAA GAA AGC CAA ACC |
| c | H1 | CAT CTC GGT TTG GCT TTC TTG TTA CCC AGG TAA CAA GAA AGC CAA ACC /3DTPA/ |
| d | H2 | TAA CAA GAA AGC CAA ACC GAG ATG GGT TTG GCT TTC TTG TTA CCT GGG /3BiodT/ |
| e | Thrombin apt1 | /5DTPA/ CAC TGG TAG GTT GGT GTG GTT GGG GCC AGT G |
| f | Thrombin apt2 | /5Biosg/ AGT CCG TGG TAG GGC AGG TTG GGG TGA CT |
| g | DNA1 | /5ThioMC6-D/ AGC CTA ATG TGC CCT TTC CA |
| h | DNA1rc | /5Biosg/ TGG AAC GCG CAA ATC AGG CT |
| i | ATP initiator | CCC AGG TAA CAA GAA AGC CAA ACC TCT TGT TAC CTG GGG GAG TAT TGC GGA GGA AGG T |
| j | Cortisol apt | GAA AAT ACA ACA AGA AAA AAC TCT CGG GAC GAC TAG CGT ATG CGC CAG AAG TAT ACG AGG ATA GTC GTC CC |
| k | Cortisol cs | TCC CGA GAC CCA GGT AAC AAG AAA GCC AAA CCT TTT CTT GTT GTA TTT TC |

Table.S1 DNA sequences in this study. /3ThioMC3-D/ refers to 3' Thiol Modifier C3 S-S. /3DTPA/ refers to 3' Dithiol modification. /3BiodT/ refers to 3' Biotin dT modification. /5Biosg/ refers to 5' Biotin modification.

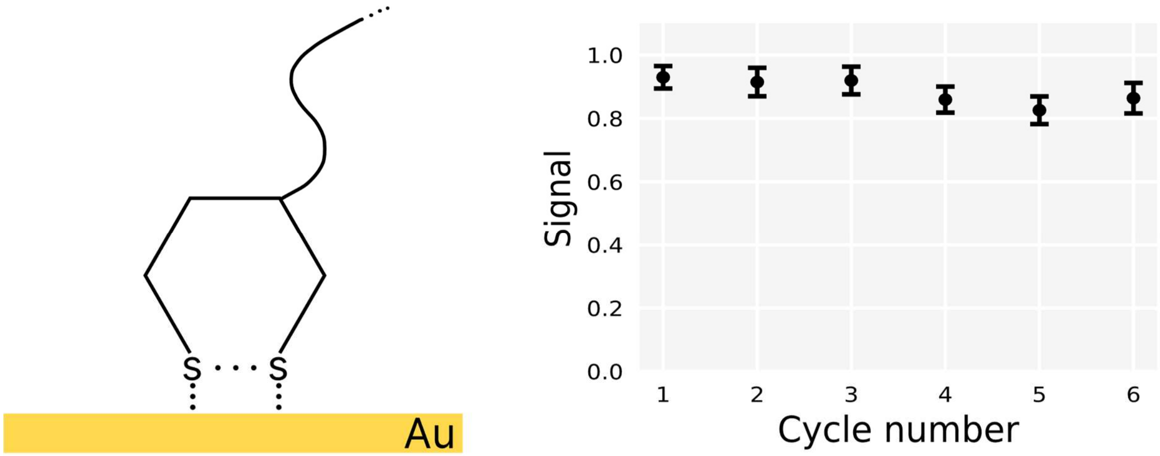


Fig.S1 Immobilization of DNA reagents with DTPA (dithiol phosphoramidite) modifications remain stable after multiple PTR cycles with minimal fluorescence decrease when testing with the DNA beacon.

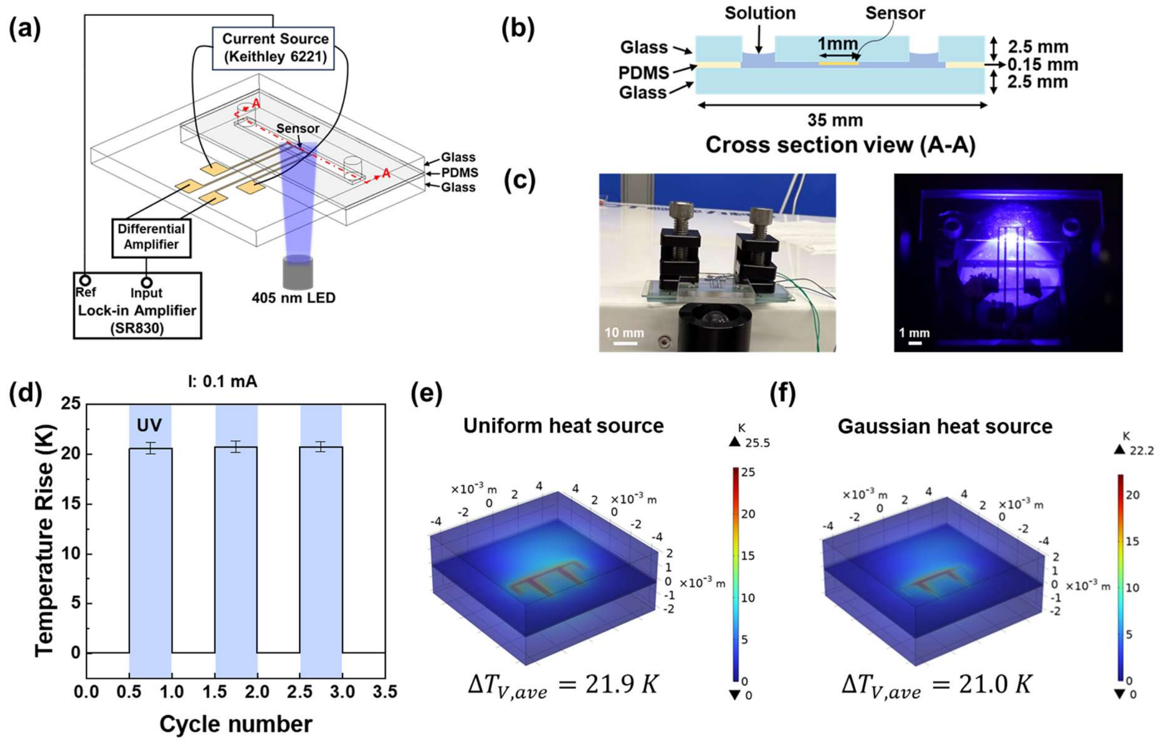


Fig.S2 The temperature increase at the gold sensor surface is measured (a-d) and modeled (e-f). The experiment and simulation results are consistent with each other and indicate a 21.9K temperature increase at the sensor surface when it is illuminated with a 405nm laser at 13608 mW/cm$^2$ power density. Two laser profiles, tophat (e) and gaussian (f), are simulated and show similar temperature increase.

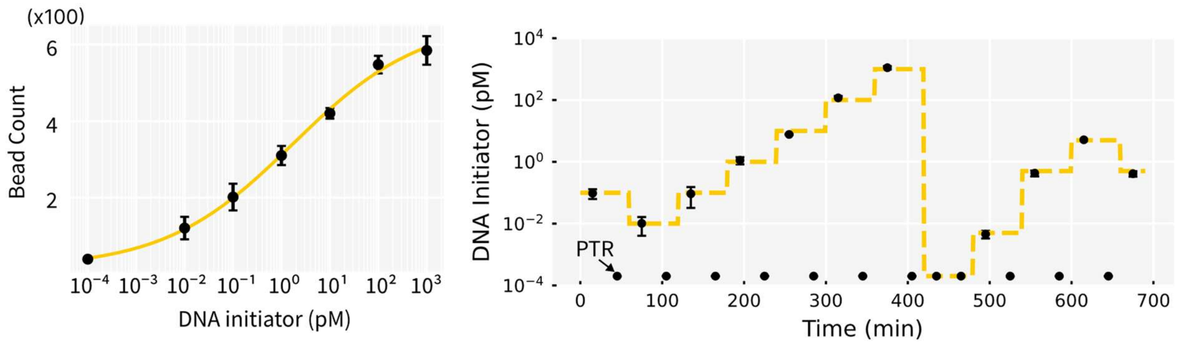


Fig.S3 Standard binding curve and continuous detection of DNA initiator strand in 1X PBS buffer. Negative control (blank) is assigned a pseudo low-concentration (0.1 aM) to fit into the binding curve plot.

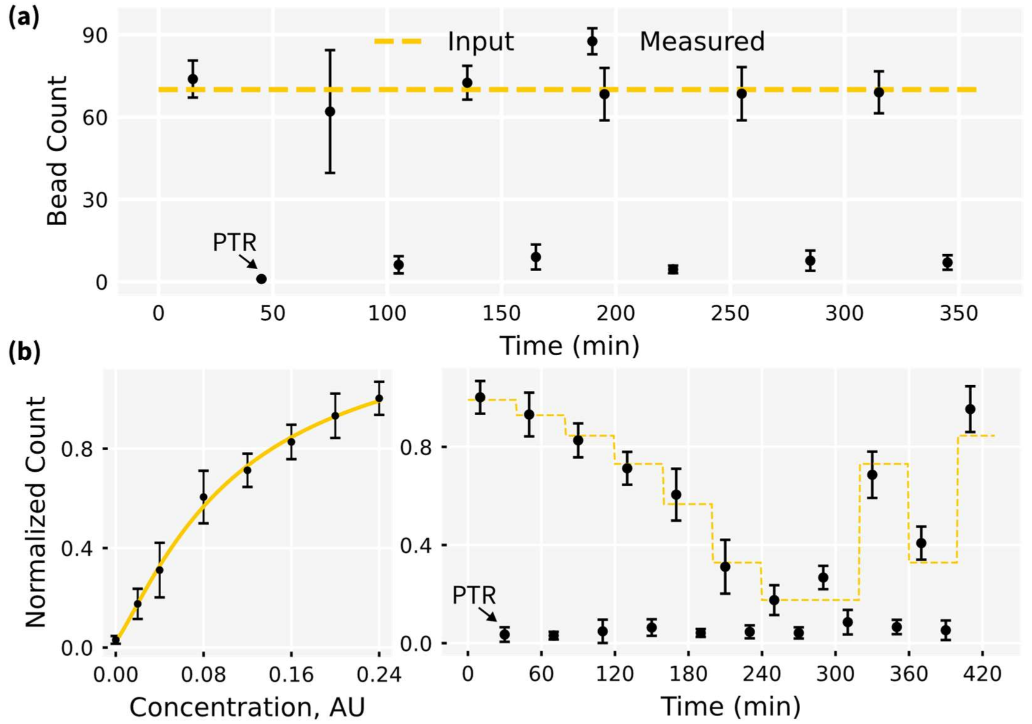


Fig.S4 DNA bead cycling with single concentration (a) and multiple concentration (b). A standard curve can also be generated with bead count against bead concentrations.

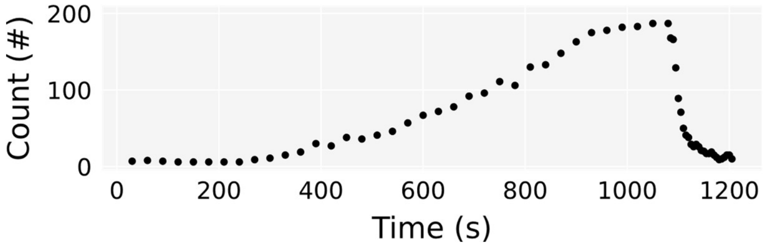


Fig.S5 Dynamic change of bead count within one measurement cycle. The number of counts gradually increase as the bead form sandwich structures on the surface with the analyte. The count suddenly decreases as the surface is illuminated with 405nm laser which heat up the sensor surface. The time step in binding phase is 30s while the time step in dissociation phase is 5s. A video showing the dynamic binding-recycling process corresponding to this plot is provided as supplement video S1.

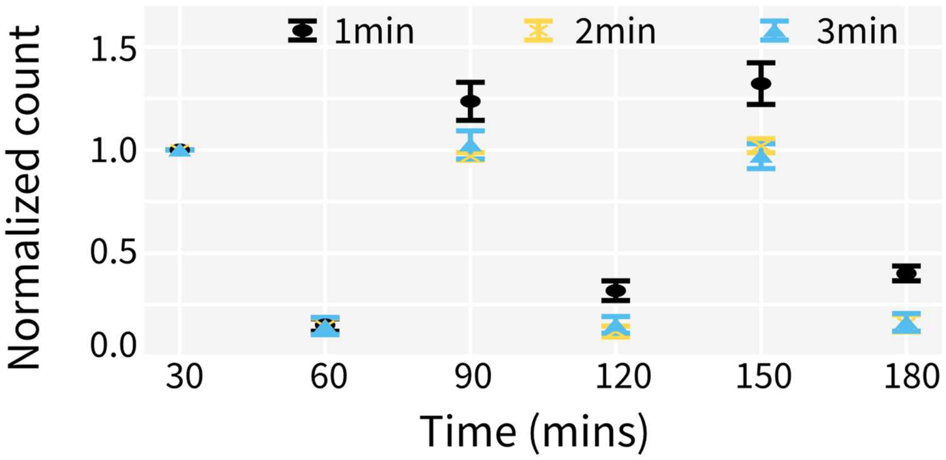


Fig.S6 PTR regeneration effect regards the laser illumination time with the 405nm laser at 13608 mW/cm$^2$ power density. PTR for 1min is insufficient to regenerate surface while 2/3mins can clean the surface well without significant damage.

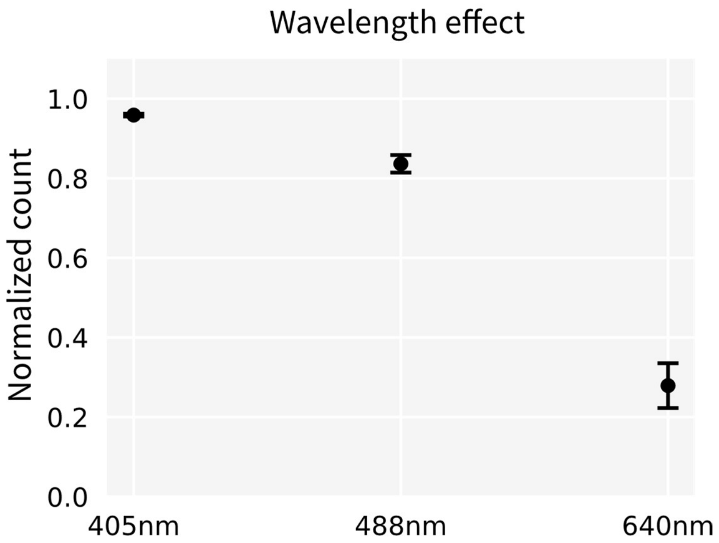


Fig.S7 PTR lifting efficiency regarding the laser wavelength. The number of lifted beads is normalized to the total number of beads and plotted as the y axis. Both 405nm and 488nm lasers can effectively remove surface attached fluorospheres at high illumination intensity. However, under the same laser intensity and photothermal heating length, 640nm only removes a small percent of beads from surface.

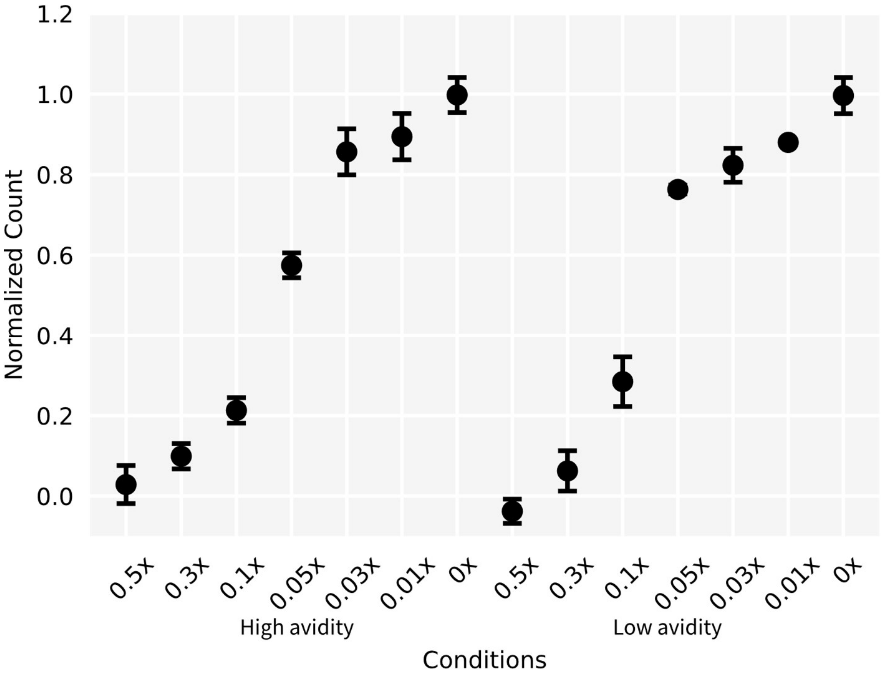


Fig.S8 The effect of avidity and ionic strength on the binding strength and PTR lifting efficiency. The amount of lifted beads are normalized to the total number of beads and plotted as the y axis. High and low avidity is achieved by adjusting the concentration of DNA1rc when labeling the avidin fluorosphere (FS). The PTR lifting efficiency of these labeled FS is tested in PBS buffer solution with different dilution factor for different ionic concentrations. As avidity increases and/or buffer ionic strength increase, the bead binds tighter to the surface and thus the PTR lifting efficiency is lower. Please see the detailed information in the Method section.

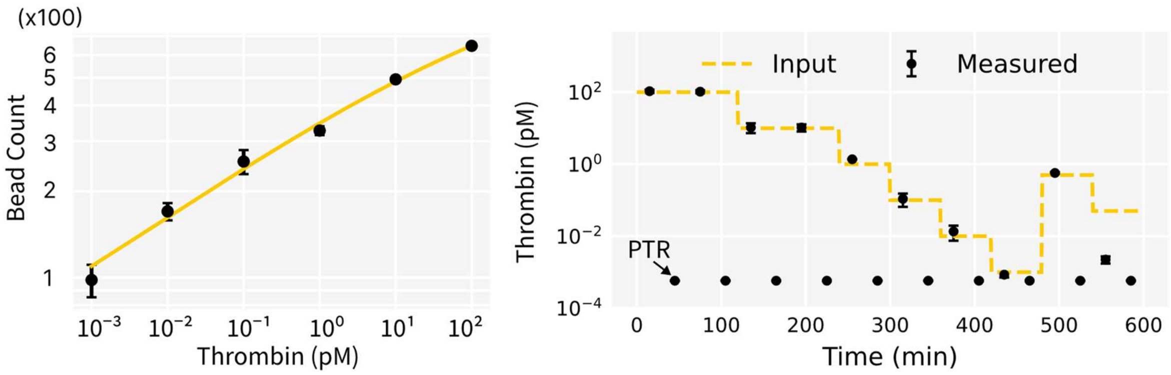


Fig.S9 PTR biosensor enables continuous ultrasensitive detection of thrombin protein in 1X PBS buffer. The thrombin binding curve is shown at the left and continuous measurement data from spike-in experiment is shown at the right. Experimental details are described in the Methods section of the main text.

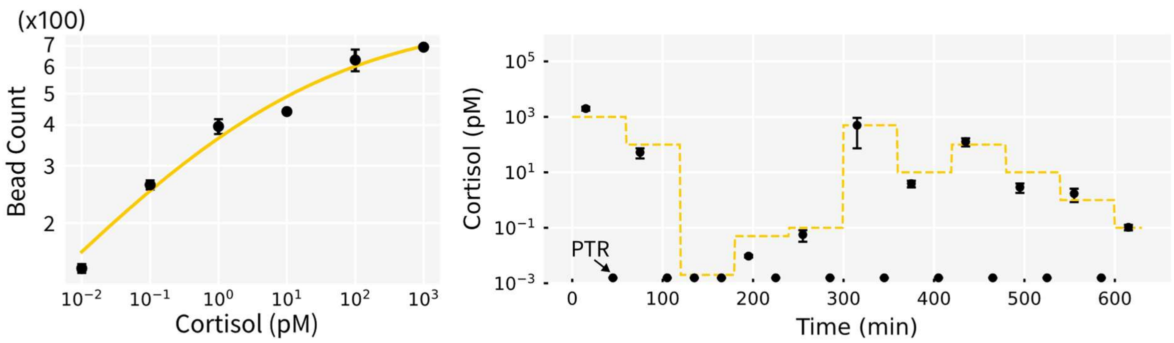


Fig.S10 PTR biosensor enables continuous ultrasensitive detection of cortisol in 1X PBS buffer. The cortisol binding curve is shown at the left and continuous measurement data from spike-in experiment is shown at the right. Experimental details are described in the Method section.

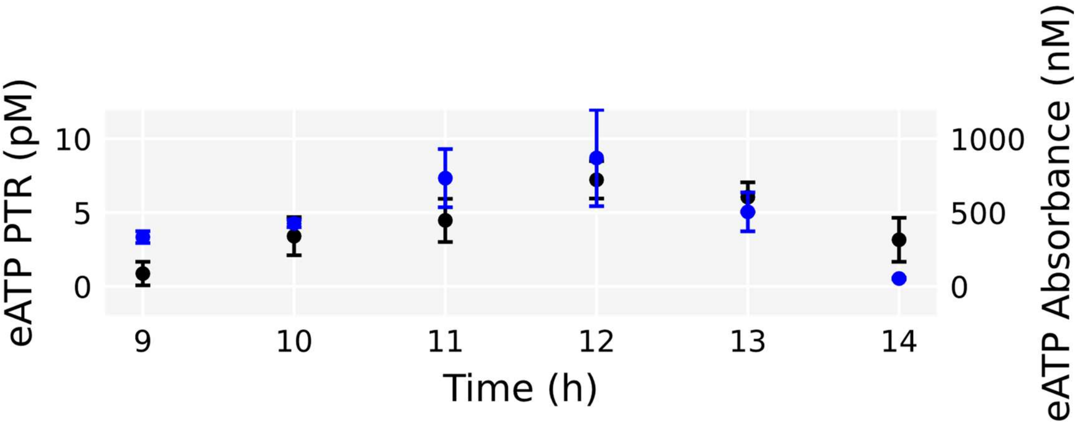


Fig.S11 PTR biosensor measures the extracellular ATP trend in E. coli. culture media (black) in the inline monitoring setup from Fig.5 in the main text. The trend matches with that measured with a commercial ATP luciferase assay (blue). Due to the high sensitivity, the PTR assay is used with sample diluted 100,000 times and only require several ul of sample, while the commercial assay typically require 200-250 ul of sample.

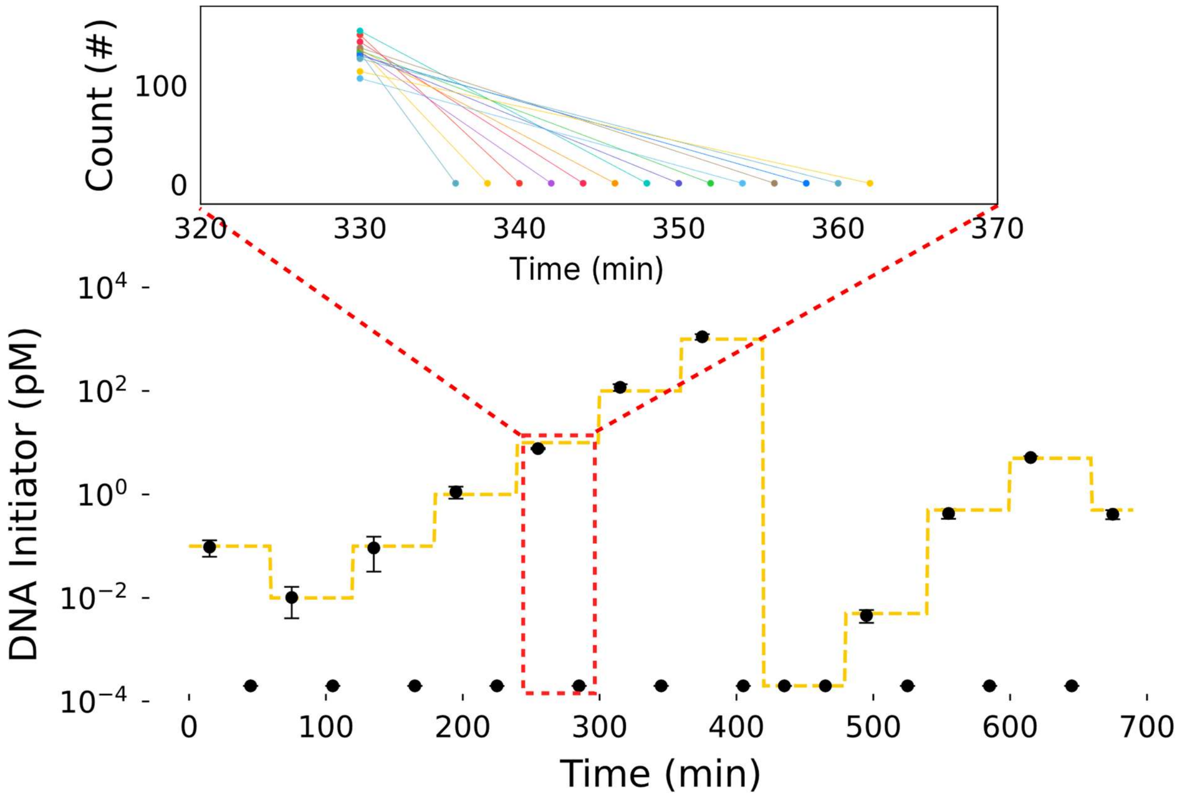


Fig.S12 Schematic to illustrate how PTR data is generated using automated microscope. For each measurement data point shown in the time-series plot, approximately 15 locations are measured, recycle sequentially and bead count averaged to produce the mean bead counts at the time interval. The sequential measurement due to limited laser illumination field, though 2 minutes of photo-thermal heating is sufficient to recycle each position.
